# Plasticity in the glucagon interactome reveals novel proteins that regulate glucagon secretion in αTC1-6 cells

**DOI:** 10.1101/373118

**Authors:** Farzad Asadi, Savita Dhanvantari

## Abstract

Glucagon is stored within secretory granules of pancreatic alpha cells until a stimulus, such as a change in microenvironmental conditions, triggers its release. We hypothesized that the secretory response of the alpha cell to various stimuli could be determined by plasticity in the network of proteins that interact with glucagon within alpha cell secretory granules. To answer this question, we isolated secretory granules from alpha TC1-6 cells and identified glucagon-interacting proteins by affinity purification coupled with liquid chromatography/tandem mass spectrometry. Proteomic analyses revealed a network of cytoplasmic and histone proteins. Specifically, the interaction between glucagon and histone H4 and the ER stress protein GRP78 was confirmed through co-immunoprecipitation of secretory granule lysates, and co-localization within secretory granules using high-resolution confocal microscopy. The composition of these networks was altered at different glucose levels (25 mM vs 5.5 mM) and in response to the paracrine inhibitors of glucagon secretion, GABA and insulin. Finally, siRNA-mediated silencing of a subset of nonhistone proteins revealed novel proteins that may regulate glucagon secretion. We have therefore described a novel and dynamic glucagon interactome within alpha cell secretory granules, and suggest that plasticity in the interactome governs the alpha cell secretory response to paracrine and nutritional stimuli.

## Introduction

In pancreatic alpha cells, glucagon secretion is governed by nutritional, hormonal and neural effectors. In the normal physiological state, this stimulus-secretion coupling is tightly regulated to maintain glucose homeostasis. However, in the case of diabetes, this tight coupling is disrupted, ^1^resulting in dysfunctional glucagon secretion, and may actually be a factor in development of type 2 diabetes ^2^. This abnormal glucagon secretion has led to strategies to control glucagon action to ameliorate the hyperglycemia of diabetes ^3^, ^4^. This strategy, although effective in the short term, tends to increase alpha cell mass and worsen alpha cell dysfunction over the long term. Therefore, a preferable strategy may be to control the secretion, rather than the action, of glucagon for improved glycemic control in diabetes.

In the context of the pancreatic islet, there is some debate as to whether glucagon secretion is primarily regulated by the paracrine influence of the beta cell, or through intrinsic factors ^5^. Both insulin and GABA secreted from the beta cell strongly inhibit glucagon secretion as does somatostatin ^6^ ^7^. However, these actions are dependent on prevailing glucose concentrations; at 5 mM glucose, glucagon secretion is maximally stimulated, but insulin secretion is not affected ^5^, suggesting that intrinsic factors may exert an equally prominent influence on glucagon secretion.

Some proposed mechanisms of intrinsic regulation of glucagon secretion include glucose metabolic-induced changes in Ca^2+^ and K^+^ membrane conductances or intracellular Ca^2+^ oscillations ^9^ ^10^. In addition to intracellular ion flux, intrinsic factors can also include proteins involved in the intracellular trafficking of glucagon. We have previously shown that prolonged culture of alpha TC1-6 cells in medium containing 25 mM glucose resulted in the up-regulation of components of the regulated secretory pathway ^11^, notably proteins that are present in or on secretory granules, such as SNARE exocytotic proteins and granins. There may be direct interactions between granule proteins, such as chromogranin A and carboxypeptidase E, to ensure proper trafficking of glucagon into secretory granules ^12^, and distinct sorting signals within glucagon may mediate these interactions^13^. Therefore, proteins within the alpha cell secretory granules that directly interact with glucagon may provide additional clues to how glucagon secretion is regulated.

To this end, we have developed a strategy using a tagged glucagon fusion protein to analyze the proteome of secretory granules isolated from alpha TC1-6 cells and identify proteins that specifically interact with glucagon. We show that this novel glucagon “interactome” exhibits plasticity in response to glucose, insulin and GABA, and have identified some novel glucagon-interacting proteins that regulate glucagon secretion.

## Results

Our method for purification of proteins that associate with glucagon within the alpha cell secretory granules consisted of two sequential steps. First, we modified and applied a previously published method ^14^ for enrichment of the secretory granule fraction. Second, we used Fc-glucagon for affinity purification to pull down proteins associated with glucagon within the secretory granules.

### Secretory granule enrichment

We have shown the enrichment of secretory granules by immunoblotting for organelle-specific markers (Supplementary Fig. 1). The final granule fraction was positive for only the secretory granule marker, vesicle associated membrane protein 2 (VAMP2) (Supplementary Fig. 1A). In contrast, the granule fraction did not contain the trans-Golgi marker TGN46 (Supplementary Fig. 1B), the nuclear envelope marker LaminB1(Supplementary Fig. 1C), or the endoplasmic reticulum marker Calreticulin (Supplementary Fig. 1D). As a positive control, the general cell lysate contained all 4 markers.

### Confirmation of the enriched secretory granules

Secretory granules in alpha cells have been previously studied using transmission electron microscopy and their average sizes have been reported to be in the range of 180-240 nm ^15^ ^16^ ^17^. We instead confirmed the presence of secretory granules using nano-scale flow cytometry with Fc-glucagon as exclusive marker for alpha cell secretory granules ^18^ ^19^. We used beads in the range of 110-880 nm for calibration in the range of the reported sizes for secretory granules (Supplementary Fig. 2A). Fc-glucagon^+^ secretory granules distributed mostly to the gated regions of 179 and 235 nm (Supplementary Figs. 2B and 2C), confirming our method of isolating secretory granules from αTC1-6 cells.

#### Proteomic analysis of proteins that are associated with glucagon within alpha cell secretory granules

The Fc or Fc-glucagon was purified from the granule lysate by affinity purification, and proteins that interact with either Fc alone or Fc-glucagon were identified with LC-MS/MS. Proteins that were pulled down by Fc alone in both 25 mM glucose (Supplementary Table S1) and 5.5 mM glucose (Supplementary Table S2) conditions were subtracted from the list of proteins identified using Fc-glucagon, thus identifying proteins that specifically interact with glucagon, which we term the” glucagon interactome”. Proteins were assigned the following categories: metabolic-secretory-regulatory, histones, cytoskeletal, and ribosomal. We identified 42 and 96 glucagon interacting proteins within the category of metabolic-regulatory-secretory proteins when the cells were cultured in media containing 25 mM (Fig. 1A) and 5.5 mM glucose (Fig. 1B), respectively.

**Figure 1.**
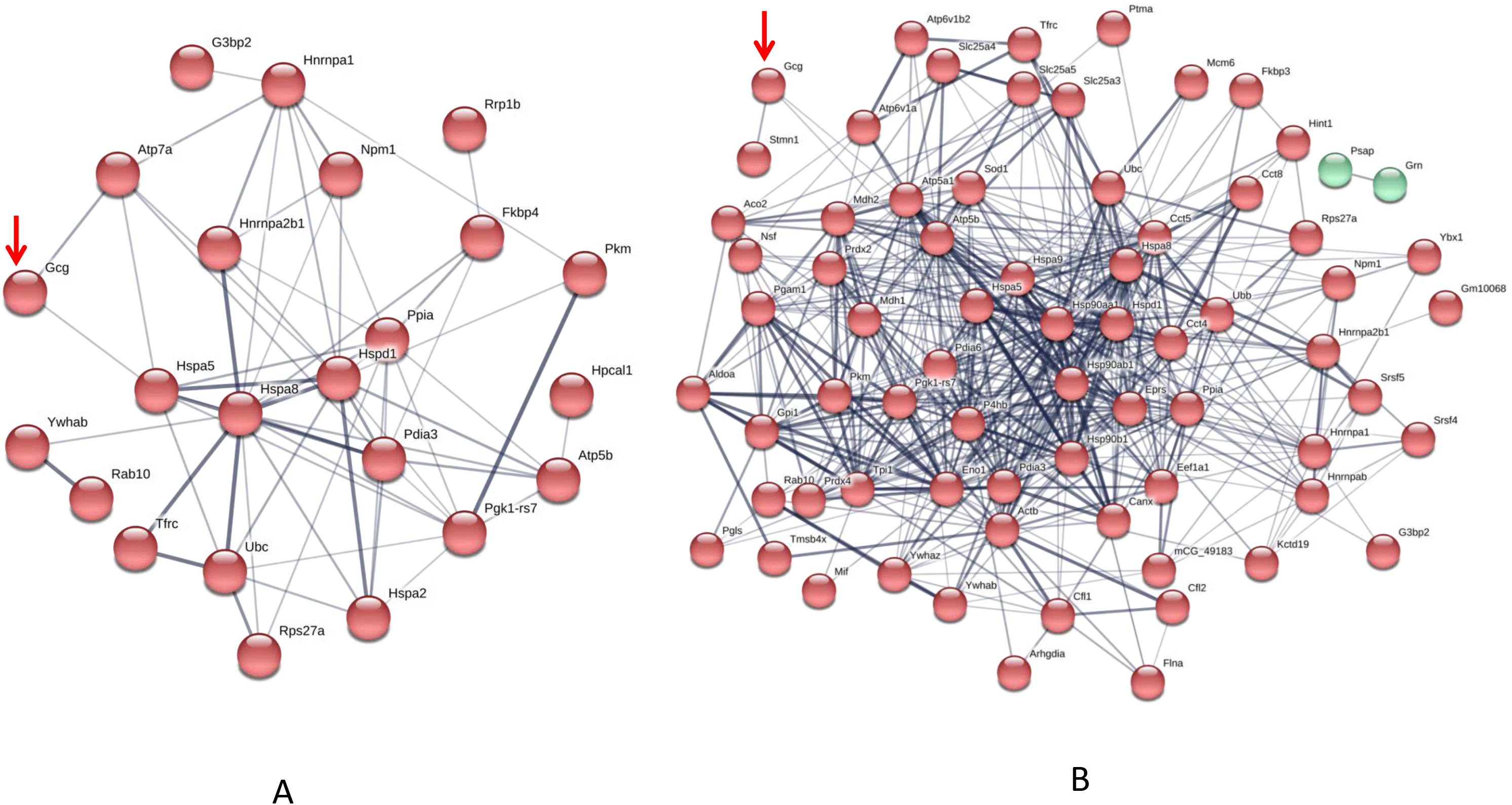
The glucagon interactome in secretory granules of αTC1-6 cells. Cells were transfected with Fc-glucagon or Fc alone, and cultured in DMEM containing 25 mM or 5.5 mM glucose for 24h. Fc-glucagon was purified from enriched secretory granules and associated proteins were identified by LC-MS/MS. **(A)** Proteomic map of the metabolic-regulatory-secretory proteins that are predicted to associate with glucagon in the context of 25 mM glucose. Network clustering predicts direct interactions between glucagon and glucose regulated protein 78 KDa (Hspa5, also known as Grp78), and ATPase copper transporting alpha polypeptide (Atp7). **(B)** Proteomic map of the metabolic-regulatory-secretory proteins that are predicted to associate with glucagon in the context of 5.5 mM glucose. Network clustering predicts direct interactions between glucagon and GRP78, stathmin1 (Stmn1), and heat shock protein 90-alpha (Hsp90aa1).

In media containing 25 mM glucose, there was a predicted direct interaction of glucagon with glucose regulated protein 78 kDa (GRP78 or Hspa5), and ATPase copper transporting alpha polypeptide (Atp7a) (Fig. 1A), while in media containing 5.5 mM glucose, GRP78, Stathmin1 (Stmn1), and Heat shock protein 90-alpha (Hsp90aa1) were predicted to directly interact with glucagon (Fig. 1B). Under conditions of either 25 mM or 5.5 mM glucose, one common predicted interaction was that between glucagon and GRP78.

#### GRP78 interacts with glucagon and co-localizes to glucagon-positive secretory granules

Affinity purification of Fc-glucagon or Fc alone from the secretory granule lysate was followed by immunoblotting for GRP78 (Fig. 2A). The presence of GRP78 immunoreactivity with Fc-glucagon, and not Fc alone, demonstrates a direct interaction with glucagon in the enriched secretory granules (Fig. 2A). Immunofluorescence microscopy showed co-localization of GRP78 and endogenous glucagon within the secretory granules in αTC1-6 cells (Fig. 2B). There was a strong positive correlation between glucagon and GRP78 immunoreactivities (PCC = 0.85 ± 0.08), indicating significant co-localization of GRP78 and glucagon.

**Figure 2:**
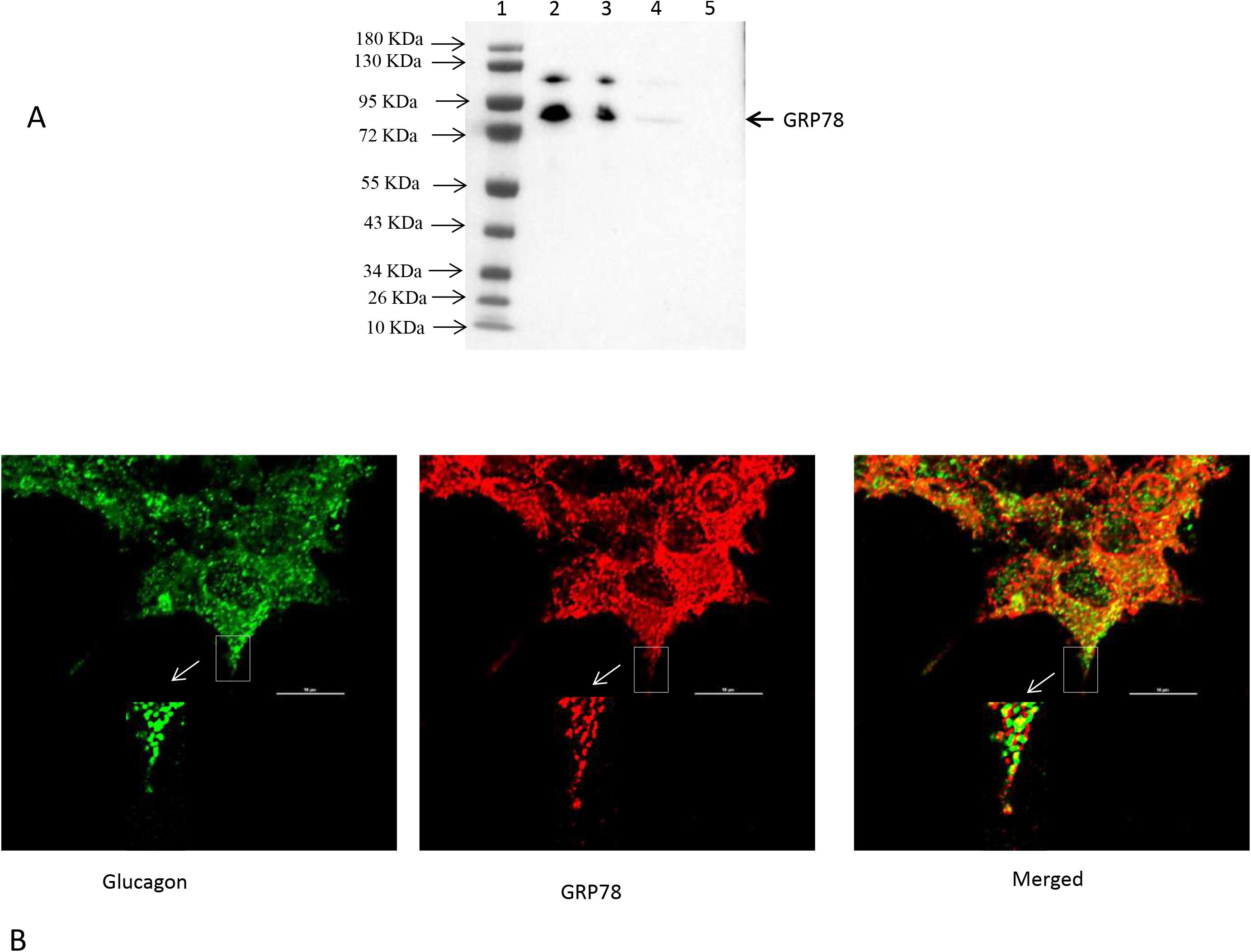
Glucagon and GRP78 directly interact and are localized within secretory granules in αTC1-6 cells. **(A)** Western blot showing GRP78 immunoreactivity in: total cell extracts from untransfected (lane 2) and transfected (lane 3) cells; affinity-purified Fc-glucagon from isolated secretory granules (lane 4); and affinity-purified Fc alone from isolated secretory granules (lane 5). GRP78 binds to Fc-glucagon, but not Fc alone. **(B)** Immunofluorescence microscopy of glucagon (green), GRP78 (red) and both images merged. Cells were cultured on collagen-coated coverslips for 24h in DMEM containing 25 mM glucose. Images were acquired and analyzed with NIS software (Nikon, Canada). Pearson correlation coefficient (PCC) indicates strong correlation between GRP78 and glucagon (PCC = 0.85 ± 0.08). ROI shows areas of colocalization of GRP78 and glucagon within secretory granules.

#### GABA induces histone H4 interaction with glucagon and presence in secretory granules

Interestingly, proteomic analysis also revealed the presence of histone proteins, along with structural proteins and ribosomal proteins, within the secretory granules in αTC1-6 cells (Supplementary Table S3). Histone H4 was predicted to interact with glucagon in cells incubated in medium containing 5.5 mM glucose. Therefore, we reasoned that this interaction was responsive to external effectors. We treated αTC1-6 cells with GABA, a well-known modulator of glucagon secretion ^20^ and examined the interaction between histone and glucagon. Co-immunoprecipitation of granule lysates, histone H4 ELISA of granule lysates, and immunofluorescence microscopy all validated the interaction of histone H4 with glucagon and presence of histone H4 in secretory granules of alpha TC1-6 cells after treatment with GABA (Figs 3). Affinity purification of Fc-glucagon or Fc alone from the secretory granule lysate was followed by immunoblotting for histone H4 (Fig. 3A). The presence of histone H4 immunoreactivity with Fc-glucagon, and not Fc alone, demonstrates a direct interaction with glucagon in the enriched secretory granules (Fig. 3A). We then confirmed the presence of histone H4 in secretory granules isolated from αTC1-6 cells by ELISA (Fig. 3B). In cells treated with GABA in 25 mM glucose, there was a detectable amount of histone H4 in the granules.

**Figure 3:**
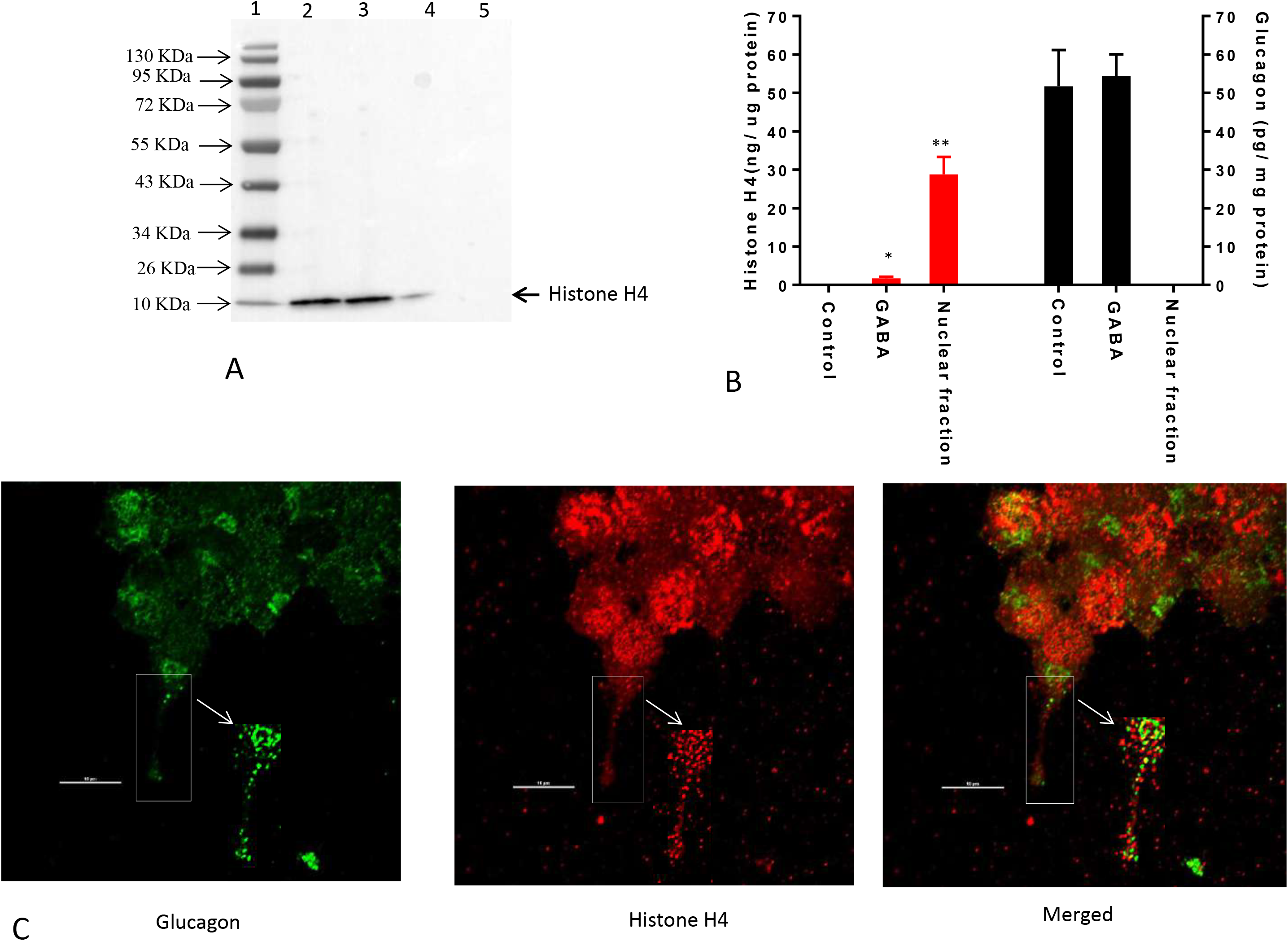
GABA induces direct interaction between glucagon and histone H4 within secretory granules in αTC1-6 cells. **(A)** Western blot shows histone H4 immunoreactivity in: total cell extracts from untransfected (lane 2) and transfected (lane 3) cells; affinity-purified Fc-glucagon from isolated secretory granules (lane 4); and affinity-purified Fc alone from isolated secretory granules (lane 5). Histone H4 binds to Fc-glucagon, but not Fc alone. **(B)** Quantitative ELISA measurement of histone H4 (left Y axis) and glucagon (right Y axis) within the secretory granules (control GABA, insulin) and the nuclear fraction of αTC1-6 cells. Values are expressed as mean±SD and compared with 1-way ANOVA (α=0.05).*p<0.05; **p<0.001. **(C)** Immunofluorescence microscopy of glucagon (green), histone H4 (red) and both images merged. Cells were cultured on collagen-coated coverslips for 24h in DMEM containing 25 mM glucose. Images were acquired and analyzed with NIS software (Nikon, Canada). Pearson correlation coefficient (PCC) indicates strong correlation between histone H4 and glucagon (PCC = 0.78 ± 0.08). ROI shows areas of colocalization of histone H4 and glucagon within secretory granules.

That this result was not due to contamination from the nuclear fraction was shown by the finding that histone H4 levels were undetectable in the secretory granules of cells not treated with GABA. As a positive control, the nuclear fraction showed high levels of histone H4. Finally, immunofluorescence microscopy showed the presence of histone H4 in glucagon-containing secretory granules (Fig. 3C), and there was significant co-localization with glucagon as assessed by Pearson’s correlation coefficient (PCC =0.78 ± 0.08).

#### The glucagon interactome changes in response to glucose, GABA and insulin

Since the interaction between histone H4 and glucagon was dependent on glucose levels and GABA, we determined the effects of the major alpha cell paracrine effectors, GABA and insulin, on the glucagon interactome. The profiles of the metabolic-regulatory-secretory proteins that associate with glucagon within secretory granules were altered upon treatment with GABA, insulin or GABA+insulin, respectively, when αTC1-6 cells were cultured in medium containing 25 mM glucose (Fig. 4) and in 5.5 mM glucose (Fig. 5). In addition, we tabulated the profiles of the histone, cytoskeletal and ribosomal proteins in response to GABA, insulin and GABA+insulin in 25 mM glucose (Supplementary Tables S4-A-C) or 5.5 mM glucose (Supplementary Tables S5-A-C). Our glucagon interactomes were functionally classified into the following groups: Binding, Structural molecule, Catalytic, Receptor, Translation regulator, Transporter, Signal transducer, Antioxidant. The proportion of proteins in each category is shown in the context of 25 mM glucose (Supplementary Table S6) and 5.5 mM glucose (Supplementary Table S7).

**Figure 4:**
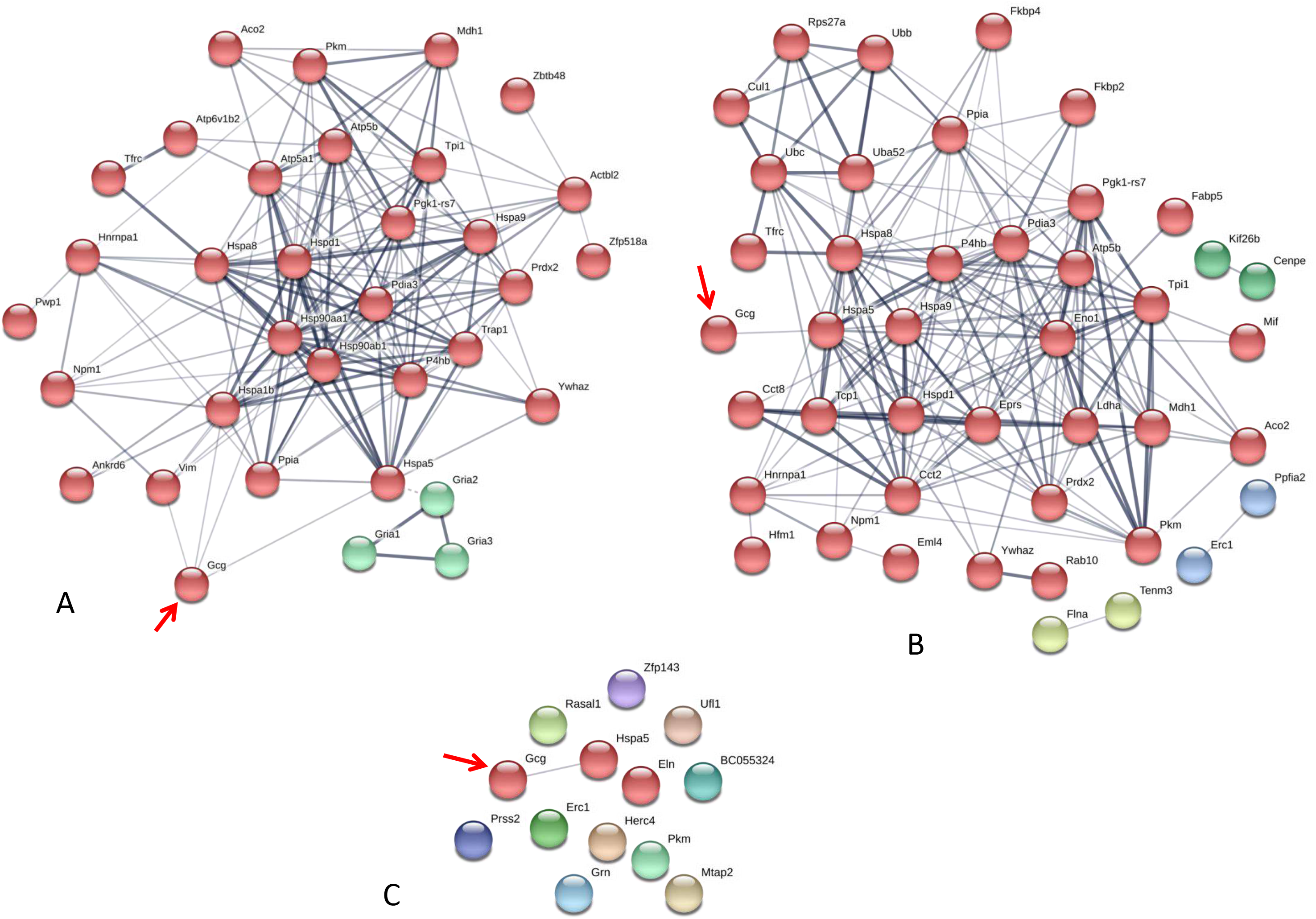
The glucagon interactome is altered in response to paracrine effectors in 25 mM glucose. αTC1-6 cells were transfected with Fc-glucagon or Fc alone, and treated with GABA (25 μM), insulin (100 pM) or GABA (25 μM) plus insulin (100 pM) for 24h in DMEM containing 25 mM glucose. Fc-glucagon was purified from isolated secretory granules and associated proteins were identified by LC-MS/MS. **(A)** Proteomic map of metabolic-regulatory-secretory proteins that are associated with glucagon after treatment of αTC1-6 cells with GABA shows direct interactions with 4 proteins: GRP78, Heat shock 70 kDa protein 1B (Hspa1b) Heat shock protein 90-alpha (Hsp90aa1), and Vimentin (Vim). **(B)** After treatment with insulin or **(C)** GABA+Insulin, glucagon is predicted to interact only with GRP78.

**Figure 5:**
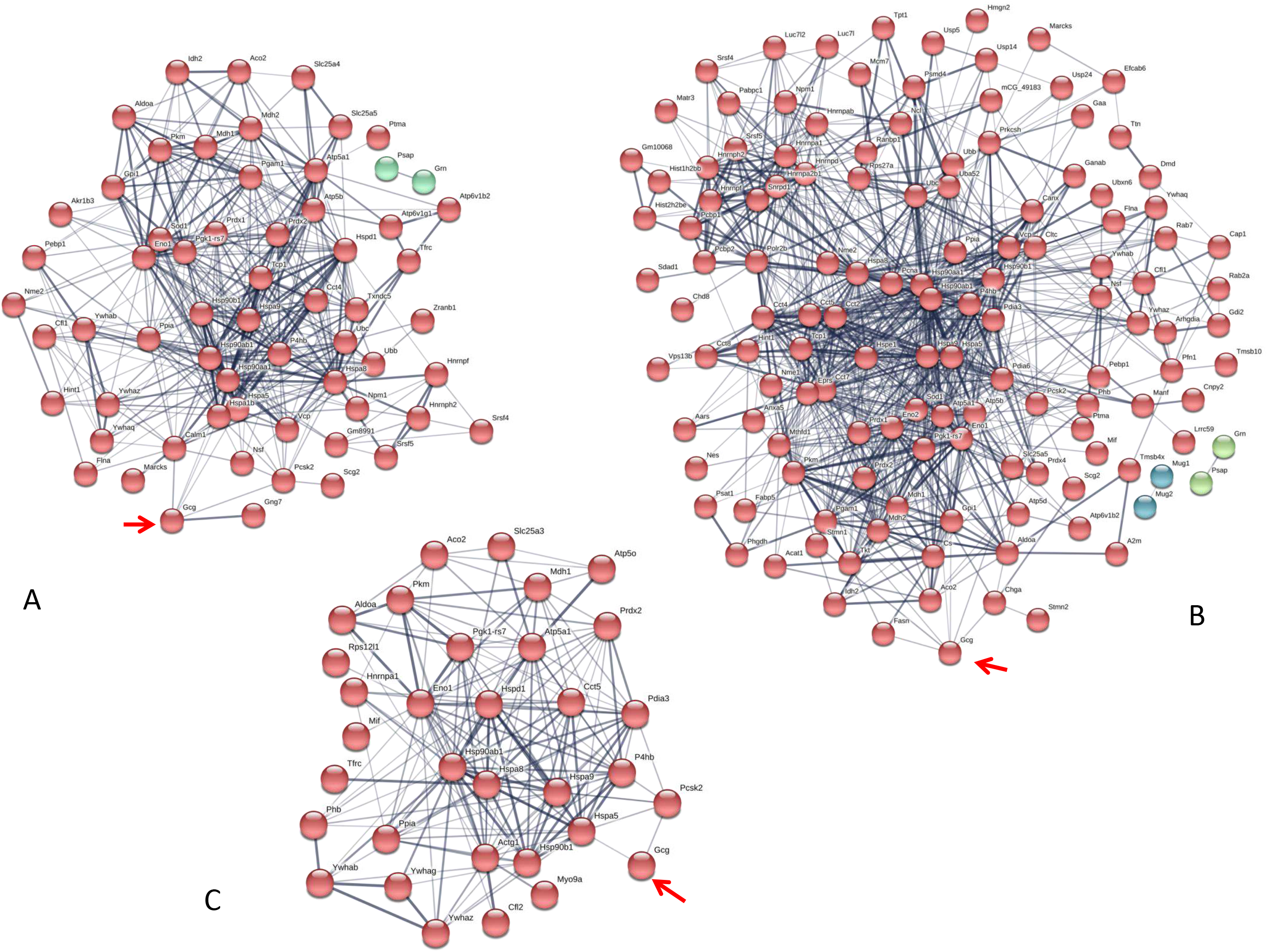
The glucagon interactome is altered in response to paracrine effectors in 5.5 mM glucose. αTC1-6 cells were transfected with Fc-glucagon or Fc alone, and treated with GABA (25 μM), insulin (100 pM) or GABA (25 μM) plus insulin (100 pM) for 24h in DMEM containing 5.5 mM glucose. Fc-glucagon was purified from isolated secretory granules and associated proteins were identified by LC-MS/MS. **(A)** Proteomic map of metabolic-regulatory-secretory proteins that are associated with glucagon after treatment of αTC1-6 cells with GABA shows direct interactions with 6 proteins: GRP78, Heat shock protein 90-alpha (Hsp90aa1), Protein convertase subtilisin/kexin type2 (PCSK2), Heat shock 70 kDa protein 1B (Hspa1b), Calmodulin 1 (Calm1), Guanine nucleotide-binding protein G(I)/G(S)/G(O) subunit gamma-7 (Gng7). **(B)** After treatment with insulin, glucagon is predicted to directly interact with 7 proteins: GRP78, Heat shock protein 90-alpha, Annexin A5 (Anxa5), Stathmin1 (Stmn1), PCSK2, Fatty acid synthase (Fasn), and Chromogranin A (Chga). **(C)** After treatment with GABA+Insulin, glucagon is predicted to directly interact with GRP78 and PCSK2.

The networks of proteins that are predicted to interact with glucagon within the secretory granules under conditions of 25 mM glucose are illustrated in Fig. 4. In cells treated with GABA, glucagon is predicted to directly interact with GRP78, HSP1B, HSP90, and vimentin (Fig. 4A); in cells treated with insulin and GABA+insulin, glucagon interacts directly with only GRP78 (Figs 4B - C). The clusters of metabolic-secretory-regulatory proteins that make up the rest of the glucagon interactomes change in composition in response to the different treatments. The numbers of proteins categorized as “structural molecule activities” decreased in response to GABA (~ 45%) or insulin (~ 38%) and increased in the GABA + insulin group (~ 16%) `compared to the control (Supplementary Table S6). The numbers of cytoskeletal proteins increased in the GABA (29%), insulin (12%) and GABA + insulin (35%) groups, while the numbers of ribosomal proteins decreased in those groups by 51%, 14% and 66%, respectively (Table 1A).

**Table 1A:** Sub-groups of proteins categorized as “structural molecules” in the glucagon interactome under conditions of 25 mM glucose. Panther GO-Slim Molecular Function analysis resulted in 3 sub-categories. The values represent protein hits as a percentage of the total number of hits within each sub-category when αTC1-6 cells were cultured in media containing 25 mM glucose.

|  | Structural constituent of cytoskeleton | Structural constituent of ribosome | Extracellular matrix structural constituent |
| --- | --- | --- | --- |
| Control | 66.7 | 29.2 | 4.2 |
| GABA | 85.7 | 14.3 | - |
| Insulin | 75 | 25 | - |
| GABA+Insulin | 90 | 10 | - |

Compared to cells incubated in medium containing 25 mM glucose, there were dramatic increases in the numbers of metabolic-regulatory-secretory proteins associated with glucagon after treatment with GABA, insulin or GABA+insulin in cells incubated in media containing 5.5 mM glucose (Fig 5). In cells treated with GABA, glucagon is predicted to directly interact with the following proteins: GRP78 (Hspa5), HSP 90alpha (Hsp90aa1), proprotein convertase subtilisin/kexin type 2 (PCSK2), heat shock 70 kDa protein 1B (Hsp1b), calmodulin 1(Calm1), and guanine nucleotide-binding protein G(I)/G(S)/G(O) subunit gamma-7 (Gng7) (Fig. 5A). Under insulin treatment, the following proteins were predicted to directly interact with glucagon: GRP78, HSP 90-alpha, annexin A5 (Anxa5), stathmin1 (Stmn1), fatty acid synthase (Fasn) and chromogranin A (Chga) (Fig. 5B); and only 2 proteins, GRP78 and PCSK2, were predicted to directly interact with glucagon after treatment with GABA + insulin (Fig. 5C).

In the context of 5.5 mM glucose, the number of cytoskeletal proteins decreased, and the number of ribosomal proteins increased compared to cells treated with GABA, insulin and GABA + insulin in 25 mM glucose (Table 1B). Interestingly, the total numbers of proteins classified as “structural molecule activities” did not change appreciably across treatments (Supplementary Table S7). However, differences became apparent when cytoskeletal and ribosomal proteins were compared separately. When compared to 5.5 mM glucose alone, there were decreases of ~24% and ~ 35%, respectively, in the numbers of cytoskeletal proteins when cells were treated with GABA or insulin alone, but a ~71% increase in response to GABA+Insulin. Conversely, the numbers of ribosomal proteins increased by ~26% and ~ 43% in response to GABA and insulin, respectively, and decreased by ~ 69% in response to GABA+Insulin (Table 1B).

**Table 1B:** Sub-groups of proteins categorized as “structural molecules” in the glucagon interactome under conditions of 5.5 mM glucose. Panther GO-Slim Molecular Function analysis resulted in 3 sub-categories. The values represent protein hits as a percentage of the total number of hits within each sub-category when αTC1-6 cells were cultured in media containing 5.5 mM glucose.

|  | Structural constituent of cytoskeleton | Structural constituent of ribosome | Extracellular matrix structural constituent |
| --- | --- | --- | --- |
| Control | 50 | 45.5 | 4.5 |
| GABA | 38.1 | 57.1 | 4.8 |
| Insulin | 32.4 | 64.9 | 2.7 |
| GABA+Insulin | 85.7 | 14.3 | - |

#### The dynamic glucagon interactome reveals novel proteins that regulate glucagon secretion

From our glucagon interactomes, we identified 11 proteins that interact with glucagon after treatment of αTC1-6 cells with either GABA or insulin in media containing 25 mM glucose. To determine their effects on alpha cell function, these proteins were depleted with siRNAs and glucagon secretion and cell content were measured. Of these 11 proteins, knockdown of ELKS/Rab6-interacting/CAST family member 1 (ERC1) increased glucagon secretion (p<0.001), while gene silencing of 14-3-3 zeta/delta (KCIP-1), cytosolic malate dehydrogenase (MDH1), FXYD domain-containing ion transport regulator 2 (FXYD2) and protein disulfide-isomerase (PDI) reduced glucagon secretion to the same statistically significant level (p<0.001). As well, knockdown of peroxiredoxin-2 (PRDX2), ATP synthase F1 subunit alpha (ATP5F1A), histone H4 and aconitate hydratase mitochondrial (ACO2) reduced glucagon secretion (p<0.01), as did knockdown of alpha-tubulin 2 (AT2) (p<0.05) (Fig. 6A). Gene silencing of MDH1, PRDX2, ATP5F1A and FXYD2 reduced cellular glucagon content to a significance level of p<0.001. Gene silencing of KCIP-1, ACO2, Histone H4 and PDI all reduced the levels of cellular glucagon content to a significance level of p<0.01 and that for ERC1 at p<0.05 (Fig. 6B). Gene silencing of GRP78 had no effect on glucagon secretion, and reduced cellular glucagon content (p<0.05).

**Figure 6:**
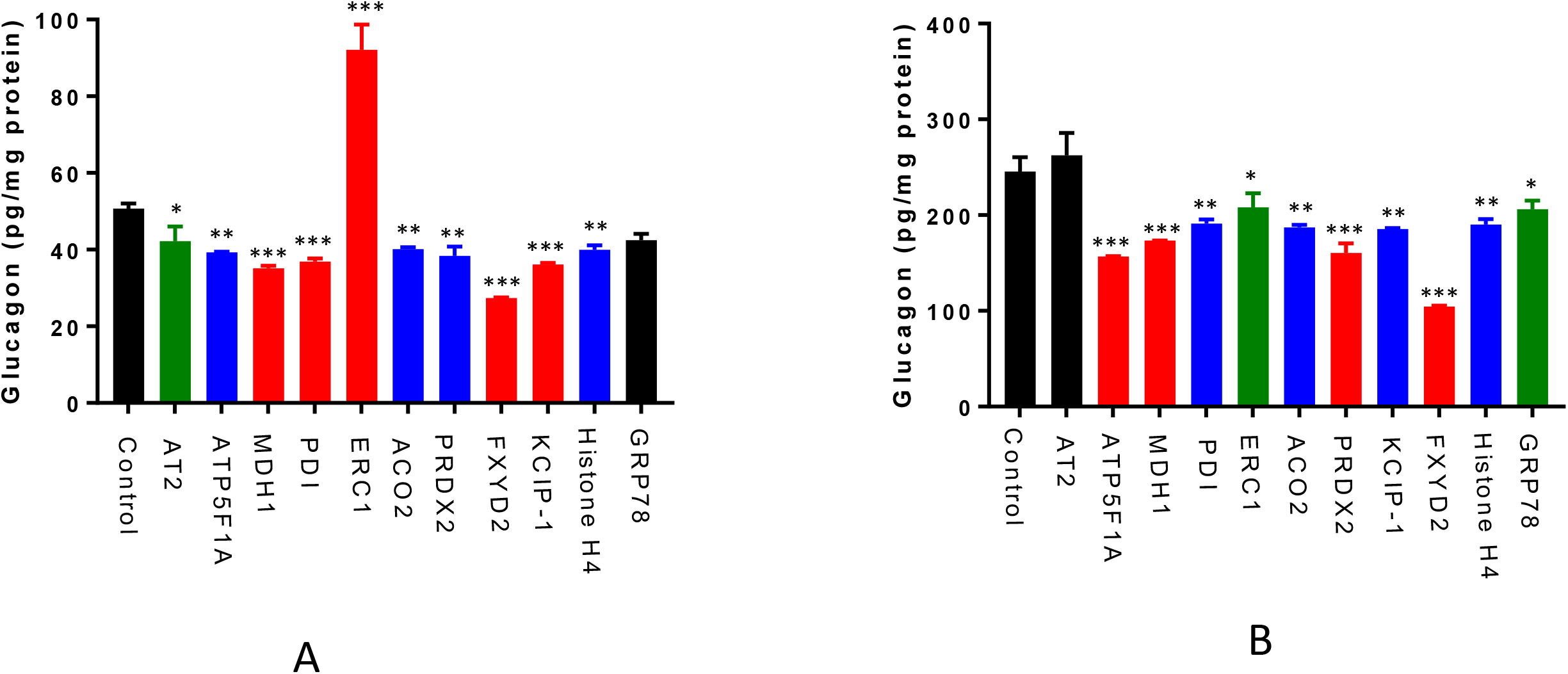
Glucagon secretion and cell content are regulated by a subset of interactome proteins. **(A)** Glucagon secretion and **(B)** cell content were assessed after siRNA-mediated gene silencing of the following proteins: Alpha-tubulin 2 (AT2), ATP synthase F1 subunit alpha (ATP5F1A), Malate dehydrogenase 1 (MDH1), Protein disulfide-isomerase (PDI), ELKS/Rab6-interacting/CAST family member 1 (ERC1), Aconitate hydratase mitochondrial (ACO2), Peroxiredoxin-2 (PRDX2), 14-3-3 protein zeta/delta (KCIP-1), FXYD domain-containing ion transport regulator 2 (FXYD2), histone H4, and GRP78 using pre-designed siRNAs for the mouse genome. After siRNA transfection, αTC1-6 cells were cultured in DMEM containing 25 mM glucose for 24h and glucagon levels were measured using ELISA. Values are expressed as mean ± SD (α=0.05). *p<0.05; **p<0.01; ***p<0.001.

## Discussion

We have identified a dynamic “glucagon interactome” within secretory granules of alpha cells that is altered in response to ambient glucose levels and the paracrine effectors GABA and insulin. To this end, we used a tagged glucagon construct, Fc-glucagon, to bring down proteins within secretory granules. We validated enrichment of the secretory granules by nano-scale flow cytometry and immunoblotting with compartment-specific markers. We identified a network of 392 proteins within the secretory granules that interact with glucagon and showed a direct interaction with GRP78 and Histone H4. Additionally, we showed that components of the interactome played a role in glucagon secretion, thus revealing a role for the interactome in the regulation of glucagon secretion.

In response to glucose concentrations alone, we identified a common backbone of 25 metabolic-regulatory-secretory proteins. The dramatic increase of 71 additional proteins when the cells cultured in 5.5 mM glucose may indicate an overall increase in protein synthesis, reflected by up-regulation in RNA-binding proteins that modulate biosynthesis of proteins that comprise the islet secretory granules ^21^. The up-regulation in chaperonin levels may also indicate an increase in protein synthesis, in accordance with its role in insulin biosynthesis ^22^. Chaperonins, as key components of the cellular chaperone machinery, are involved in maturation of newly-synthesized proteins in an ATP dependent manner ^23^. As ATP-generating proteins, such as ATP5F1A, MDH1 and glucose metabolic proteins, were also increased, we speculate that 5.5 mM glucose induced a stress response that results in increased protein translation. This hypothesis is strengthened by the identification of cold shock protein, peroxiredoxin, thiol-disulfide isomerase and thioredoxin within the glucagon interactome at 5.5 mM glucose, all of which are up-regulated in pancreatic islets in response to stress ^22^.

One protein that was consistently predicted as interacting directly with glucagon was the ER stress protein and molecular chaperone GRP78. Previous proteomic studies have identified GRP78 in islets and beta cells ^24^ ^22^. Its presence in alpha cell secretory granules may not be surprising, as it has previously been found in non-ER compartments such as the nucleus and lysosomes. Our data suggest that GRP78 may be a novel sorting receptor for glucagon in the regulated secretory pathway of alpha cells. We have previously shown a potential role of chromogranin A as a sorting receptor for glucagon in both αTC1-6 cells and PC12 cells^13^, but unlike GRP78, we did not demonstrate any direct interactions with glucagon. While knockdown of GRP78 did not reduce glucagon secretion, it did reduce cell content, indicating a potential role in intracellular trafficking, but not exocytosis, of glucagon.

Interestingly, we identified histone proteins as a functional part of the glucagon interactome. The discovery of histone proteins within alpha cell secretory granules is novel, and supported by the findings that the cytosolic fraction of pooled islets from multiple human donors had abundant amounts of the histone H2A ^25^, which is also the most abundant histone in the glucagon interactome. As well, quantitative proteomics of both αTC1 and ßTC3 cells revealed the presence of histones H4, H3, H2A, H2B and H1 ^26^. For the first time, we show that one of these histones, H4, directly binds to glucagon and may regulate its secretion. It has been suggested that histones contained within secretory granules in neutrophils could function as a defence mechanism, whereby histones H2A and H2B interact with the plasma membrane to generate extracellular traps in response to bacterial infections ^27^. Thus, it is possible that histone proteins in the glucagon interactome take a role in the fusion step of granule exocytosis. Additionally, secretion of histones and other nuclear proteins has been associated with an inflammatory or senescent secretory phenotype ^28^ ^29^.

The alpha cell paracrine effectors, GABA and insulin, remodeled the glucagon interactome in αTC1-6 cells in a manner that was dependent on the ambient glucose levels. Upon GABA treatment, the number of proteins in the glucagon interactome increased by 25 and 73 unique proteins in the context of 25 mM and 5.5 mM glucose, respectively. Compared to the respective control groups, GABA altered >70% and >80% of the metabolic-regulatory-secretory proteins within the glucagon interactome in the context of 25 mM and 5.5 mM glucose, respectively. One potentially novel GABA-regulated protein that may function in glucagon secretion in 25 mM glucose is ERC1, which has a role in the formation of the cytomatrix active zone and insulin exocytosis from beta cells ^30^, and we show for the first time a potential inhibitory effect of ERC1 on glucagon secretion that may be dependent on GABA. Another potentially novel player in GABA-regulated glucagon secretion is KCIP-1, associated with beta cell survival ^31^. Furthermore, our proteomics findings suggest that GABA may enhance glucose uptake and glucose tolerance through leucine-rich repeat proteins. These proteins bind to the insulin receptor to promote glucose uptake in beta cells ^32^, and thus may be a new paracrine, or even autocrine, regulator of alpha cell function. Interestingly, in the context of 5.5 mM glucose, GABA recruited PCSK2 and secretogranin 2, known alpha cell granule proteins that function in proglucagon processing^13^. Although our previous work showed no changes in PCSK2 in response to 5.5 mM glucose ^11^, we now show that plasticity in PCSK2 expression may be due to GABA under these glucose concentrations.

Treatment with insulin remodeled the glucagon interactome with 36 common proteins and 38 unique proteins in 25 mM glucose and 142 unique proteins in 5.5 mM glucose. In the context of 25 mM glucose, insulin treatment increased the number of biosynthetic proteins, consistent with its role in cellular growth. Kinesin-like proteins also increased, suggesting a potential role in alpha cell secretory granule synthesis and glucose homeostasis, as has been documented in beta cells ^33^. In the context of 5.5 mM glucose, insulin up-regulated nucleoside diphosphate kinases A and B, proposed regulators of insulin secretion ^34^. We also identified the small G proteins SAR1, Rab2A and RhoA, present in INS-1 cell secretory granules ^35^; however, their functions are not completely known.

Interestingly, treatment of the cells with GABA+ insulin in 25 mM glucose caused a dramatic decrease in the overall numbers of proteins within the glucagon interactome. Interaction with GRP78 remained preserved, while a new protein, microtubule-associated protein 2, appeared in the glucagon interactome. This protein may have a potential role in glucose homeostasis, as it is down-regulated in isolated diabetic rat islets exposed to low glucose conditions ^36^. In the context of 5.5 mM glucose, the combination of GABA and insulin again predicted the presence of PCSK2 in the glucagon interactome, as seen with GABA treatment alone. One intriguing hypothesis is that the interaction between glucagon and PCSK2 is stabilized under these conditions in αTC1-6 cells, and invites revisiting the question of PCSK2 acting as a sorting receptor for glucagon^13^.

In conclusion, we have described a novel and dynamic glucagon interactome that is remodeled in response to glucose and the alpha cell paracrine effectors, GABA and insulin. Our proteomics approach has revealed a number of novel secretory granule proteins that function in the regulation of glucagon secretion and illustrates the plasticity in the protein components of the alpha cell secretory granules. These findings provide an important proteomics resource for further data mining of the alpha cell secretory granules and targeting diabetes treatment.

## Materials and methods

Sources for all reagents, assays, and software packages are listed in Supplementary Table S8.

### Gene construct and plasmid preparation

We designed a glucagon fusion construct [Fc-glucagon–pcDNA3.1(+)] using the amino acid sequence of glucagon derived from human proglucagon (GenScript, USA; http://www.genscript.com) attached to the 3’ end of the cDNA encoding the CH2/CH3 domain of mouse IgG-2b (called Fc), preceded by a 28 amino acid signal peptide as described previously 24. As a negative control for all transfection, proteomics, immunofluorescence microscopy and co-immunoprecipitation experiments, we designed and used an Fc-pcDNA3.1(+) construct. The sequences of the purified Fc and Fc-glucagon constructs were confirmed at the London Regional Genomics Facility, University of Western Ontario.

### Extraction and enrichment of secretory granules

Wild type pancreatic αTC1-6 cells (a kind gift from C. Bruce Verchere, University of British Columbia, Vancouver, BC) were cultured in DMEM containing 25 mM glucose, L-glutamine, 15% horse serum and 2.5% fetal bovine serum, as described previously ^11^. The cells were grown to 90% confluency in three 10cm dishes for each experimental condition. Cells were transfected with Fc alone or Fc-glucagon using Lipofectamine 2000 in Opti-MEM (ThermoFisher Scientific) plus 10% FBS for 16h. Cells were then incubated for 24 h in DMEM containing 25 mM glucose prior to the granule enrichment procedure. To determine changes in granule size, mass and proteome, cells were incubated with or without GABA (25 μM), insulin (100 pM), or GABA (25 μM) plus insulin (100 pM) in either 25 mM or 5.5 mM glucose for 24h prior to the granule enrichment procedure. Each experimental condition was done in 3-6 biological replicates.

After the 24h incubation period, media were removed and cells were washed in ice-cold HBSS prior to processing for granule enrichment based on a previously published method ^14^ with some modifications. Briefly, cells were detached using 5mM EDTA in PBS (pH 7.4) containing Mini Protease Inhibitor Cocktail (Supplementary Table S8) on ice. Cells were collected into microfuge tubes, centrifuged at 500 × g for 5 min and resuspended in 1.2 mL ice-cold homogenization buffer (20mM Tris-HCl pH7.4, 0.5mM EDTA, 0.5mM EGTA, 250mM sucrose, 1 mM DTT, Complete Protease Inhibitor Cocktail and 5 μg/mL Aprotinin).The cells were passed 10 times through a 21G needle and again 10 times through a 25G needle. The resulting lysates were centrifuged at 900 × g for 10 min at 4°C to obtain a post-nuclear supernatant (PNS). The nuclear fraction was washed 7 times in ice-cold homogenization buffer and kept at −80°C. The PNS was centrifuged at 5400 × g for 15 min at 4°C to obtain a post-mitochondrial supernatant, which then was spun at 25000 × g for 20 min at 4°C to pellet the granule enriched fraction. The granule enriched fraction was re-suspended in 8 mL ice-cold homogenization buffer and spun at 24000×g for 20 min at 4°C. This washing step was repeated once at 24000xg, then 4 times at 23000×g, 22000×g, 21000×g, and 20000×g. Finally, the secretory granules were washed at 20000×g for 18min to obtain the final granule-enriched preparation. Enrichment was confirmed through immunoblotting for markers of the secretory granules, endoplasmic reticulum, trans-Golgi network, and nuclear membrane marker as described below.

### Immunoblotting for organelle-specific markers

The enriched preparations of secretory granules from αTC1-6 cells were lysed using non-ionic lysis buffer (50 mM Tris pH 7.4, 150mM NaCl, 1% Triton X-100 plus Complete Protease Inhibitor Cocktail and 5 μg/mL Aprotinin). Proteins were resolved on a 4-12% SDS-PAGE gel (NuPAGE), transferred to a PVDF membrane and probed with the following antibodies (Supplementary Table S8) to identify different subcellular compartments: vesicle-associated membrane protein 2 (VAMP2) for mature secretory granules; calreticulin for the endoplasmic reticulum; TGN46 for the trans-Golgi network; and Lamin B1 for the nuclear envelope. The immunoreactive bands were visualized using HRP-conjugated goat anti-rabbit secondary antibody and Clarity Western ECL substrate. Images were acquired on a BioRad ChemiDoc Imaging System. Total cell extracts were used as positive controls.

### Nanoscale flow cytometry

#### a) Secretory granule preparation

We used nano-scale flow cytometry (A50-Micro nanoscale flow cytometer; Apogee FlowSystems Inc.) to confirm enrichment of the secretory granules and to determine the size distribution of the granules. To this end, αTC1-6 cells were transfected with Fc-glucagon or Fc alone and secretory granules were extracted and enriched as described above. The extracted secretory granules were fixed in freshly prepared 2% PFA in PBS (pH 7.4) for 15 min and then permeabilized with 0.5% saponin in 1% BSA for 20 min at room temperature. The fixed and permeabilized granules were centrifuged at 25000 ×g for 20 min at 4°C and washed 3 times in 0.1% saponin in PBS. Fluorescent labelling of Fc-containing granules for nanoscale flow analysis was conducted with FITC-IgG (1:250 dilution in 0.1% saponin in 1% BSA/PBS) in the dark for 1h. The labelled secretory granules were diluted 200 times in 0.1% saponin in PBS for nanoscale flow cytometry.

#### b) Size calibration

Secretory granules of non-transfected cells were used for size calibration. To this end, ApogeeMix beads were used to establish sizing gates along the Y axis - large angle light scattering (LALS) versus X-axis-small angle light scattering (SALS) plot 29 30. The microparticle mixture contained plastic spheres of 180nm, 240nm, 300nm, 590nm, 880nm and 1300 nm diameter with refractive index 1.43 and 110nm and 500 nm green fluorescent beads with refractive index 1.59. Based on the manufacturer established default, the calibrated gates of the size distribution of the nanoparticles were 110 nm, 179 nm, 235nm, 304 nm, 585 nm, and 880 nm, and these gates were used to analyse and categorize subpopulations of the enriched secretory granules.

#### c) Nano-flow analysis

Samples of the enriched secretory granule preparations were aspirated at a flow rate of 1.5μL/min. To count the numbers of Fc-glucagon+ granules, the green fluorescence of FITC excitation (L488) was gated and the numbers of Fc-glucagon+ granules were counted at 110nm, 179nm, 235nm, 304nm, 585nm and 880nm within the LALS vs. L488 plot. In this respect, to get the LALS vs. L488 plot, its gate was normalized for the following isotypes: secretory granules of non-transfected cells, secretory granules of Fc transfected cells, FITC-IgG and diluent. This method resulted in size distribution of the granules that were positive for Fc-glucagon, specifically. All experiments were done in 3 biological samples and values were expressed by descriptive analysis as percent distribution of the gated granules.

### Proteomic analysis of secretory granule proteins associated with glucagon

#### a) Granule lysate preparation

Alpha TC1-6 cells were transfected with either Fc alone (negative control) or Fc-glucagon and treated with effectors (GABA, insulin and GABA plus insulin) in media containing 25mM or 5.5 mM glucose as described above. Secretory granules were extracted and enriched as described above, and lysed in a non-ionic lysis buffer (50 mM Tris pH 7.4, 150 mM NaCl, 1% Triton X-100). Lysates were stored at −80°C until analysis.

#### b) Affinity purification

The Fc or Fc-glucagon was purified from the granule lysate by immunoprecipitation as we have done previously ^12^. To this end, a slurry of Protein A-Sepharose beads (Supplementary Table S8) was prepared by mixing washed beads with binding buffer (50 mM Tris pH 7) at a ratio of 3:1. The granule lysate was mixed with the bead slurry in a ratio of 1:1 and rotated overnight at 4°C. The mixture was then centrifuged at 500× g for 2 min at 4°C and the pellet was washed twice with 50mM Tris (pH 7.5) and once with pre-urea wash buffer (50 mM Tris pH 8.5, 1 mM EGTA, 75 mM KCl). Then, Fc or Fc-glucagon was eluted from the beads with 2 volumes of urea elution buffer (7 M urea in 20 mM Tris buffer pH 7.5 plus 100 mM NaCl). This step was repeated twice more and the supernatants were collected and pooled. The pooled supernatant was mixed with acetone in a 1:4 ratio and kept at −20°C overnight. Then, the mixture was centrifuged at 16000×g for 15 min at 4°C and the pellet was air dried for proteomic analysis.

#### c) Proteomic analysis

Identification of proteins that associated with glucagon was conducted using LC-MS/MS according to the protocols of the University of Western Ontario Mass Spectrometry Laboratory (Dr. Don Rix Protein Identification Facility, Siebens-Drake Research Institute, London, ON, Canada; (http://www.biochem.uwo.ca/wits/bmsl/protocols.html). Briefly, the air-dried pellet was reconstituted in 50 mM NH4CO3, and proteins were reduced in 200 mM dithiothreitol (DTT), alkylated in freshly prepared 1M iodoacetamide and digested with trypsin (Sequencing Grade Modified trypsin; Promega) for 18h at 37°C with occasional shaking. The tryptic peptides were acidified using formic acid (0.25; v/v) loaded onto a Hypersep C18 column, washed, and eluted in 50% acetonitrile. The eluent was dried down in a speed vacuum and reconstituted in acetonitrile for further analysis. Each experimental condition was done in 3 biological replicates.

Peptide sequences were identified using the mouse database and further analyzed for protein categorization through PANTHER GO term analysis (www.Pantherdb.org), functional protein-protein interaction clustering through http://string-db.org and determination of subcellular locations and activity using www.uniport.org.

Proteins identified using Fc alone as bait were subtracted from the proteins pulled down by Fc-glucagon to obtain the profile of proteins that specifically interact with glucagon.

### Immunoprecipitation-immunoblotting of proteins associated with glucagon

To validate the interaction between GRP78 and glucagon or histone H4 and glucagon within secretory granules, we purified Fc-glucagon or Fc from the secretory granule preparation, followed by immunoblotting for GRP78 or histone H4. To this end, the secretory granule lysate was incubated with Protein A-Sepharose beads overnight at 4°C with rotation. The Fc or Fc-glucagon complex was eluted from the beads with 0.1 M glycine buffer (pH 2.8). The eluate was concentrated 50 times using a speed vac, run on a 10% Bis-Tris NuPAGE gel (Supplementary Table S8) and proteins were transferred onto a PVDF membrane. After an overnight incubation with primary antibodies against GRP78 or histone H4 (Supplementary Table S8), bands were visualized using HRP-conjugated goat anti-rabbit secondary antibody and Clarity Western ECL substrate. Images were acquired on a BioRad ChemiDoc Imaging System.

### Histone H4 assay

Levels of histone H4 in the secretory granule fraction were measured using a colorimetric assay. To this end, enriched secretory granule fractions were prepared, resuspended in 0.2 N HCl, passed 10 times through a 30G needle, and kept at 4°C overnight. The reaction was stopped by addition of 0.2 volumes of 1N NaOH. The supernatant was collected after centrifugation at 6500 × g at 4°C for 10 min. Protein levels were determined by BCA microplate assay, and 100 ng of protein was used for measuring total histone H4 (Histone H4 Modification Multiplex Assay Kit, Supplementary Table S8), as per the manufacturer’s instructions. The nuclear fraction was also assayed for histone H4 as a positive control.

### Immunofluorescence microscopy

To validate the presence of GRP78 and histone H4 in glucagon-positive secretory granules, alpha TC1-6 cells were cultured on collagen 1-coated coverslips, and processed for immunofluorescence microscopy as described previously ^13^ after 24h incubation in DMEM containing 25 mM glucose. Briefly, the cells fixed in 4% paraformaldehyde and permeabilized in 0.1% saponin in 0.5% BSA for 1h. After blocking in 10% goat serum in 1% BSA, Cells were then incubated with the primary antibodies (mouse anti-glucagon and rabbit anti-GRP78 or rabbit anti-histone H4, Supplementary Table S8) overnight. Then, coverslips were washed in PBS and incubated with goat anti-mouse Alexa Fluor IgG 488 and goat anti-rabbit Alexa Fluor 594 (Supplementary Table S8) for 3h in the dark at room temperature. The coverslips were mounted using ProLong Gold Antifade Mountant. Images were acquired on a Nikon A1R Confocal Microscope (Nikon, Mississauga, ON, Canada) with a × 60 Nikon Plan-Apochromat oil differential interference contrast objective lens using the NIS software (Nikon, Canada).

Analysis was done using NIS software through Pearson correlation coefficient, PCC as we described previously^13^.To this end, we prepared three coverslips per transfection and imaged them for analysis. Image analysis was performed by NIS software (Nikon, Canada), using the co-localization option. Regions of interest were manually drawn around distinct single or multicell bodies, positive for Fc-glucagon and either GRP78 or histone H4 and cropped for analysis. Co-localization of the pixels from each pseudo-colored image were used to calculate Pearson’s correlation coefficient, as we described previously^13^.

### siRNA-mediated depletion of targeted proteins

After treatment of alpha TC1-6 cells with GABA and/or insulin in media containing 25 mM glucose as described above, the proteomes were tabulated, and Venn diagram analysis revealed 27 metabolic/regulatory/secretory proteins and 36 histone/cytoskeletal/ribosomal proteins that were common between the groups treated with GABA and insulin. We selected 11 of these proteins for siRNA-mediated depletion: PRDX2, MDH1, ACO2, KCIP-1, ERC1, AT2, ATP5F1A, Histone H4, FXYD2, and PDI. siRNAs targeting each of these proteins were chosen from pre-designed siRNAs (Silencer siRNA, Thermo Fisher Scientific Inc. MA, USA (https://www.thermofisher.com/ca/en/home/life-science/rnai/synthetic-rnai-analysis.html). These proteins were chosen based on availability of the pre-designed siRNA.

Gene silencing was done based on the well-documented protocol of the Mossё et al (2000)^37^. Briefly, αTC1-6 cells were cultured to 60% confluency and transfected with final concentrations of 50 nM of pooled siRNAs (three siRNAs for each target) or control scrambled siRNA using Lipofectamine2000. After 16h, transfection media were removed and replaced with DMEM without FBS. Cells were incubated for 48h, after which media were again removed and replaced, and after 24h, expression levels of the targeted proteins were evaluated by immunoblotting using primary antibodies against each protein target as listed in Supplementary Table S8.

### Glucagon measurement

To measure cellular and secreted glucagon levels after siRNA-mediated gene silencing, cell lysates or media were acidified in HCl-ethanol (92:2 v/v) in a 1:3 ratio, kept at −20°C overnight, then centrifuged at 13000 × g for 15 min at 4°C. The supernatant was then mixed in a 1:1 ratio with 20mM Tris buffer, pH 7.5 26 and glucagon levels were measured by ELISA (ThermoFisher Scientific, Supplementary Table S8) according to the manufacturer’s instructions. Glucagon values were compared among groups by one-way ANOVA using Sigma Stat 3.5 (α = 0.05) (Systat Software Inc, Point Richmond, CA, USA).

## Data availability

All data generated or analysed during this study are included in this manuscript (and its Supplementary Information files). Raw MS/MS files are available from the corresponding author upon request.

## Acknowledgment

We would like to thank Paula Pittock at the Siebens-Drake Research Institute, University of Western Ontario for assistance in LC-MS/MS analysis. This work has been financially supported by a Discovery Grant from the Natural Sciences and Engineering Research Council of Canada to SD.

## Author contribution

Both FA and SD designed the experiments, wrote and prepared the manuscript text and figures, and reviewed the manuscript prior to submission.

